# Methanotrophic flexibility of ‘*Ca*. Methanoperedens’ and its interactions with sulfate-reducing bacteria in the sediment of meromictic Lake Cadagno

**DOI:** 10.1101/2024.11.08.622632

**Authors:** Maider J. Echeveste Medrano, Guangyi Su, Lucas A. Blattner, Pedro Leão, Irene Sánchez-Andrea, Mike S. M. Jetten, Cornelia U. Welte, Jakob Zopfi

## Abstract

The greenhouse gas methane is an important contributor to global warming, with freshwater sediments representing important potential methane sources. Anaerobic methane-oxidizing archaea mitigate methane release into the atmosphere by coupling the oxidation of methane to the reduction of extracellular electron acceptors or through interspecies electron transfer with microbial partners. Understanding their metabolic flexibility and microbial interactions is crucial to assess their role in global methane cycling. Here, we investigated anoxic sediments of the meromictic freshwater Lake Cadagno (Switzerland), where ‘*Ca*. Methanoperedens’ and sulfate-reducing bacteria co-occur, with metagenomics and long-term incubations. Incubations were performed with different electron acceptors, revealing that manganese oxides supported highest CH_4_ oxidation potential but enriched for ‘*Ca*. Methanoperedens’ phylotypes that were hardly present in the inoculum. Combining data from the inoculum and incubations, we obtained five ‘*Ca.* Methanoperedens’ genomes, each harboring different extracellular electron transfer pathways. In a reconstructed *Desulfobacterota QYQD01* genome we observed large multi-heme cytochromes, type IV pili, and a putative loss of hydrogenases, suggesting facultative syntrophic interactions with ‘*Ca.* Methanoperedens’. We also screened for putative extrachromosomal elements in the ‘*Ca*. Methanoperedens’ genomes, including BORGs. This research deepens our understanding of the metabolic flexibility and potential interspecific interactions of ‘*Ca.* Methanoperedens’ in freshwater lakes.

## Introduction

Anaerobic oxidation of methane (AOM) is an important biological sink for this potent greenhouse gas (Knittel & Boetius, 2009, Saunois *et al*., 2020, Wallenius *et al*., 2021) in a wide range of anoxic ecosystems, including inland waters, coastal and ocean ecosystem sediments (Rosentreter *et al*., 2021, Gao *et al*., 2022, Su *et al*., 2022). AOM is catalyzed via reverse methanogenesis by different groups of anaerobic methane-oxidizing (ANME) archaea (Timmers *et al*., 2017, Chadwick *et al*., 2022) using a variety of electron acceptors (Glodowska *et al*., 2022). In marine sediments, sulfate dependent-AOM (S-AOM) is the predominant process, carried out by the ANME groups 1, 2A, 2B, 2C and 3 in syntropy with sulfate-reducing bacteria (SRB) (Metcalfe *et al*., 2021, Yu *et al*., 2021, Murali *et al*., 2023). The ANME’s and SRB’s engage in direct interspecies (DIET) or extracellular electron transfer (EET) via multi-heme *c*-type cytochromes (MHC) or conductive pili and/or other intermediates (Wegener *et al*., 2015, Scheller *et al*., 2016, Krukenberg *et al*., 2018). In freshwater systems, sulfate is much less abundant and the ANME-2D group, ‘*Candidatus (Ca.)* Methanoperedens’ spp., appears to drive AOM with nitrate, humic acids, or metal oxides as electron acceptors without the need for a syntrophic partner (Haroon *et al*., 2013, Ettwig *et al*., 2016, Vaksmaa *et al*., 2017, Cai *et al*., 2018, Leu *et al*., 2020, Valenzuela *et al*., 2020, Cai *et al*., 2022, Pelsma *et al*., 2023). However, interactions with certain guilds such as nitrite-scavenging anammox or *Methylomirabilis* bacteria might be beneficial for *Methanoperedens* (Arshad *et al*., 2017, Dalcin Martins *et al*., 2022).

Reports of ‘*Ca.* Methanoperedens’ spp. in estuarine and marine habitats are rather scarce although enrichment cultures have been shown to withstand marine salinities (Frank *et al*., 2023, Echeveste Medrano *et al*., 2024a). Enrichment cultures of ‘*Ca.* Methanoperedens’ have usually been established from freshwater source material, with different electron acceptors including nitrate, manganese-, and/or iron oxides. These studies revealed that the reduction of nitrate is performed by a membrane bound nitrate reductase within the respiratory chain, while large MHC containing proteins are predicted to engage in EET for insoluble metal-oxides or electrodes (Leu *et al*., 2020, Cai *et al*., 2022, Zhang *et al*., 2023, Ouboter *et al*., 2024). As MHCs are also responsible for the electron transfer between S-AOM ANMEs and their partner SRB bacteria (Meyerdierks *et al*., 2010, McGlynn *et al*., 2015), the presence of multiple MHCs in ‘*Ca.* Methanoperedens’ spp. begs the question of whether they could also live in syntropy with SRBs in sulfate-rich freshwater ecosystems.

One of the earliest reports of the potential of ‘*Ca.* Methanoperedens’ spp. mediating freshwater S-AOM came from sediments of the meromictic Lake Cadagno (Switzerland) (Su *et al*., 2020), where anoxic incubation experiments with methane, different electron acceptors, combined with molybdate as an inhibitor of bacterial sulfate reduction, revealed that AOM was predominantly sulfate-dependent. Moreover, 16S rRNA gene-based analyses showed a striking co-occurrence along the sediment depth profile of several ‘*Ca.* Methanoperedens’ Amplicon Sequencing Variants (ASVs) and one single ‘*Desulfobulbaceae*’ ASV, now reclassified as ‘uncultured *Desulfobacterota’* according to SILVA v138.2. This observation suggests that ‘*Ca.* Methanoperedens’ is probably not responsible for the reduction of sulfate itself but might oxidize methane in syntrophy with an SRB partner.

In this study, we aim to elucidate the diversity and the metabolic potential of the ‘*Ca.* Methanoperedens’ phylotypes in the sediments of Lake Cadagno, as well as of their putative syntrophic SRB partner. For this, we combined data from long-term sediment incubations with different electron acceptors and ^13^C-methane oxidation, 16S rRNA gene amplicon sequencing and metagenomic analyses. Our findings deepen the understanding of the mechanisms involved in freshwater S-AOM, and support a potential syntrophic interaction with ‘*Ca.* Methanoperedens’ and a co-occurring *Desulfobacterota*.

## Materials and Methods

### Study workflow

In Lake Cadagno sediments, rates of anaerobic oxidation of methane (AOM-R), the abundances of ‘*Ca.* Methanoperedens’ and the uncultured *Desulfobacterota* ASVs peaked at intermediate depths (19-25 cm), where methane, sulfate, and sulfide were present at concentrations of 2-3 mM, ∼100 µM, 300-600 µM, respectively (Figure 1A). Figure 1A provides a graphical representation of the relevant data aforementioned of Figures 1, 4 and Supplementary Figure 4 from (Su *et al*., 2020). Based on these results we selected specific sediment layers for metagenomic sequencing (23 cm & 25 cm) and for further long-term slurry incubation experiments (depths 19-24 cm) (Figure 1B).

**Figure 1.**
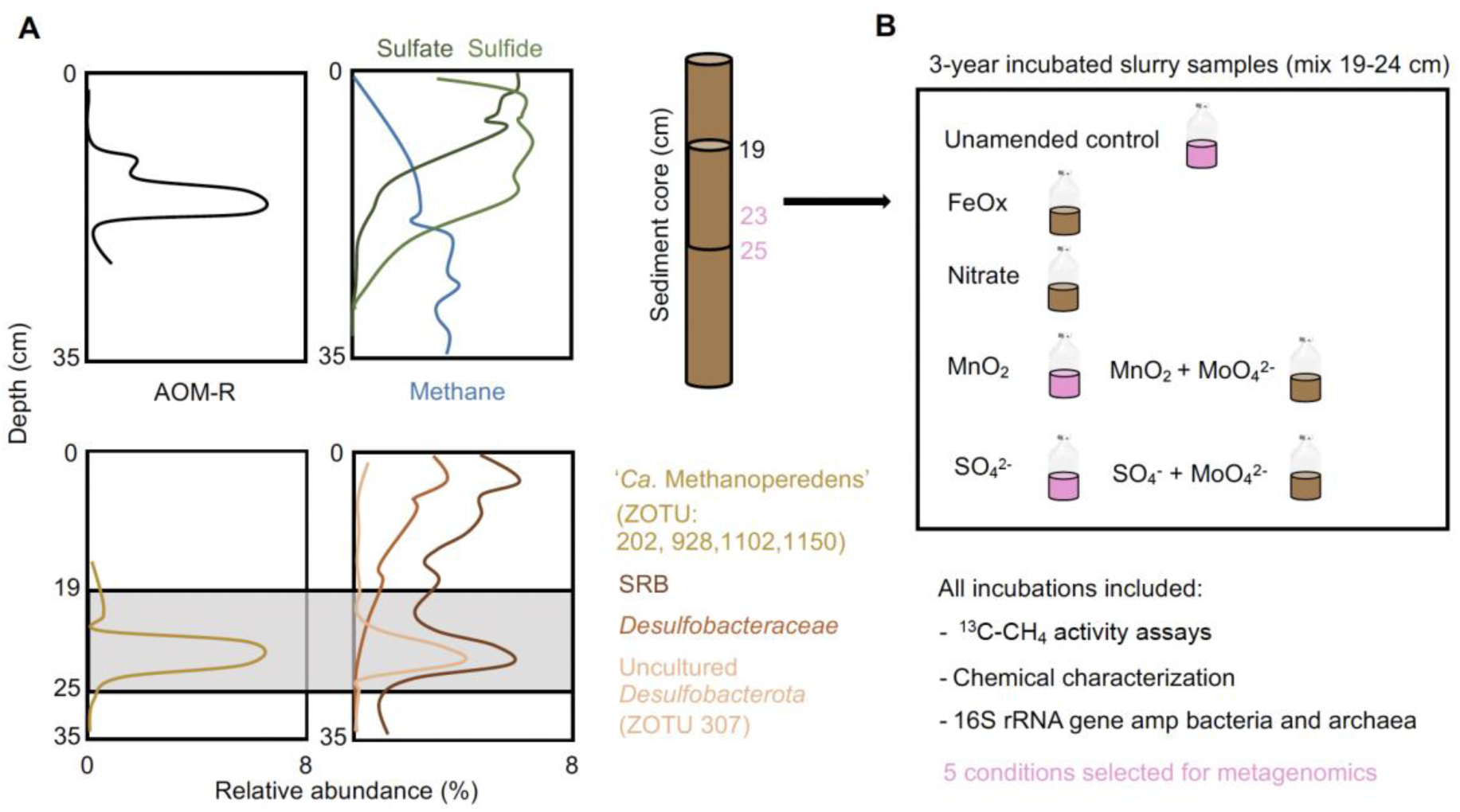
**A** Cartoon summarizing previous anaerobic oxidation of methane (AOM) data from Lake Cadagno sediment (Su *et al*., 2020). Top: Depth-specific *in situ* anaerobic oxidation of methane oxidation rates (AOM-R) determined using ^13^C-CH_4_ tracer whole-core incubations, and profiles of porewater concentrations of dissolved methane, sulfate, and sulfide. Bottom: Depth profiles of 16S rRNA gene amplicons (relative abundances) of ‘*Ca.* Methanoperedens spp.’ and SRBs in the sediment. **B** Overview of long-term slurry incubation assays presented in the current study, including the seven conditions employed and the parameters analyzed. The sediment samples and incubations selected for metagenomics are colored in pink.

### Long-term slurry incubations

We established static slurry incubations with fresh sediment material (depths 19 - 24 cm), collected from the center of Lake Cadagno (46°33’01.89’’N, 8°42’42.04E) in October 2017. In an anoxic chamber with an N_2_ atmosphere, we mixed the sediment material with anoxic freshwater medium in a volume ratio of ∼ 1:4, and dispensed 100 mL of this slurry into 250 mL serum bottles. We supplemented the slurries with either NO_3_^-^ (0.5 mM), synthesized δ-MnO_2_ (10 mM), ferrihydrite (Fe_5_O_8_H·H_2_O, 10 mM), or sulfate (4.8 mM). Molybdate (MoO_4_ ^2-^) (20 mM) was added as sulfate reduction inhibitor to incubations supplemented with MnO_2_ (10 mM) and sulfate (Fig. 1B). Finally, we purged the mixed slurries with He before injecting 20 mL ^13^C-CH_4_ (99.8 atom %, Campro Scientific). All incubation bottles were kept in the dark in an anoxic chamber at 25 °C for 36 months.

### Chemical characterization of slurry samples

At the end of the incubation, we collected the liquid phase of the slurries and filtered it through 0.45-µm filters for nutrient analysis. Sulfate was quantified by ion-chromatography (940 Professional IC Vario, Metrohm, Switzerland), while Mn^2+^, Fe^2+^, and H_2_S were determined by ICP and photometry, as described previously (Cojean *et al*., 2020). DIC concentrations were quantified on a TOC analyzer (Shimadzu, Corp., Kyoto, Japan). The carbon isotopic composition of DIC was determined by acidifying 1 mL filtered supernatant with 200 µL 85% H_3_PO_4_ in 12 mL He-flushed exetainers. The δ^13^C of the released CO_2_ was determined in the headspace after overnight equilibration via a gas-bench coupled to a Delta V Plus IRMS. The solid phase of incubation slurries was centrifuged and lyophilized prior to elemental analysis. Total carbon (TC) and total organic carbon (TOC, after acidification of the samples) as well as their δ^13^C values were determined by EA-IRMS using a Delta V Plus IRMS and a ConFlow IV interface (Thermo Fisher Scientific, Bremen Germany). Stable carbon isotope values are reported in the conventional δ-notation (in ‰) relative to the Vienna Pee Dee Belemnite standard (V-PDB). Sulfate was quantified by ion-chromatography (940 Professional IC Vario, Metrohm, Switzerland), while Mn^2+^, Fe^2+^, and H_2_S were determined by ICP and photometry, as described previously (Cojean *et al*., 2020).

### DNA extraction, amplicon sequencing and metagenomics

We extracted DNA from the inocula sediments and the incubation slurries using the FastDNA^TM^ Spin Kit for Soil (MP Biomedicals, city, country) and performed amplicon sequencing of 16S rRNA genes following the two-step PCR approach (https://support.illumina.com/documents/documentation/chemistry_documentation/16s/16s-metagenomic-library-prep-guide-15044223-b.pdf) with primers 515F-Y (5′-GTGYCAGCMGCCGCGGTAA) and 926R (5′-CCGYCAATTYMTTTRAGTTT-3’) (Parada *et al*., 2016), targeting the V4 and V5 regions of the 16S rRNA gene. Amplicons were sequenced at the Genomics Facility Basel (University of Basel/ETHZ). Details of library preparation, sequencing, quality control, and bioinformatical processing are described in (Su *et al*., 2023). We used SINTAX (v11.0.667_i86linux64, Edgar, 2016) and the SILVA 16S rRNA gene reference database v138.2 (Quast *et al*., 2013) to assign taxonomy to amplicon sequencing variants, and also to update the taxonomy of the previously described ‘*Desulfobulbaceae*’ SRB (ZOTU 307) in Su *et al*. 2020.

Furthermore, we selected two different sediment-depths (23 cm and 25 cm) and three long-term AOM incubations (unamended control, sulfate, and manganese oxides) for downstream metagenomics (Figure 1B). Library preparation and sequencing of selected sediment and incubation DNA samples was performed at the Lausanne Genomic Technologies Facility, University of Lausanne, Switzerland (https://www.unil.ch/gtf). Paired-end sequencing (150 cycles) was done on an Illumina HiSeq 2500 using the Nextera DNA Flex protocol, generating ∼ 7 Gbp/sample for all samples except for the manganese oxides. Read quality was assessed with FASTQC v0.11.9 before and after quality trimming performed with BBDuk (BBTools v38.75). Trimmed reads were co-assembled *de novo* using metaSPAdes v3.14.1 (Nurk *et al*., 2017) and mapped to assembled contigs using BBMap (BBTools v38.75) (Bushnell, 2014). Five different assemblies were generated: (i) including sediment depths 23 cm and 25 cm reads, using the reads from the control (ii), sulfate (iii), and manganese oxides (iv) slurry incubations, and (v) a bigger co-assembly including all metagenome reads (i + ii + iii+ iv) together. Contigs ≥1000-bp length were used as template for read mapping. Sequence mapping files were handled and converted using Samtools v1.10., later used for binning with CONCOCT v2.1 (Alneberg *et al*., 2014), MaxBin2 v2.2.7 (Wu *et al*., 2016), and MetaBAT2 v2.12.1 (Kang *et al*., 2019).

Generated metagenome-assembled genomes (MAGs) were dereplicated with DAS Tool v1.1.1 (Sieber *et al*., 2018) and taxonomically classified with the Genome Taxonomy Database Toolkit GTDB-Tk v2.1.0 (Chaumeil *et al*., 2019). Metagenomic mapping statistics were generated via CheckM v1.1.2 (Parks *et al*., 2015). For metagenomic binning, taxonomical read-recruitment assessment, and biogeography studies SingleM v0.16.0 and Sandpiper (https://sandpiper.qut.edu.au/) were used (May 2024), respectively (Woodcroft *et al*., 2024). MAG completeness and contamination was estimated with CheckM2 v1.0.1 (Chklovski *et al*., 2023). Open read frames (ORFs) in the MAGs were predicted and translated using Prodigal v2.6.3 (Hyatt *et al*., 2010). MAGs proteins’ were annotated with DRAM v1.5.0 (Shaffer *et al*., 2020) with default options, except min_contig_size at 1000 bp, and METABOLIC v4 (Zhou et al., 2022). Additionally, we searched for genes of interest in the annotation files via BLASTp, applying an e-value cut-off of 10^-5^. For a more accurate functional prediction of hydrogenases, we used the curated HydDB classifier http://services.birc.au.dk/hyddb/ (Søndergaard *et al*., 2016). To corroborate poorly annotated genes/proteins, manual curations were validated with the NCBI Batch Entrez Conserved Domains and InterPro (Blum *et al*., 2021). In this study, we consider microorganisms as ‘canonical SRBs’ if they encode the DsrD subunit of the dissimilatory sulfite reductase. This subunit has been identified as a marker for the enzyme’s reductive directionality, specifically for the reduction of sulfite to hydrogen sulfide (Diao *et al*., 2023). The search for proteins involved in SRB and ANME aggregate formation was done using DIAMOND with a list of protein InterPro IDs recovered from Murali et al., 2023 (IPR039662, IPR053783, IPR025295, IPR035903, PF01833) with e-value and identity cut-offs of 10^-5^ and 30%, respectively. Average amino acid or nucleotide identity (AAI or ANI) from MAGs was obtained using the FastAAI or FastANI-matrix tool option (Rodriguez-R & Konstantinidis, 2016), using 90% AAI or 95% ANI for species cutoff (Jain *et al*., 2018, Konstantinidis *et al*., 2022).

### Multi-heme c-type cytochrome (MHC) search and domain tree

We used the assembled metagenomes to investigate the presence and type of MHC on the five ‘*Ca.* Methanoperedens’ MAGs recovered together with the suspected syntrophic SRB ‘*Desulfobacterota* class QYQD01’. We first screened the selected MAGs for putative MHCs, identified by ORFs with ≥3 CXXCH motifs and used FeGenie v1.2 (Garber *et al*., 2020) annotations to differentiate Omcz nanowires, and BLASTp to identify Extracellular Cytochrome Nanowires (ECN). Subsequently, we used InterProScan v5.44-79.0 (Jones *et al*., 2014) to identify domains classified as “multi-heme cytochromes” from the identified ‘*Ca.* Methanoperedens’ MHC-harboring proteins. For this analysis, we also included reference bioreactor enrichment MAGs of ‘*Ca.* Methanoperedens’ with confirmed metatranscriptomic evidence for MHC expression under various conditions, including: *’Ca.* Methanoperedens ferrireducens’ (Fe-AOM) (Cai *et al*., 2018), *’Ca.* Methanoperedens manganicus (Mn-1)’ and *’Ca.* Methanoperedens manganireducens (Mn-2)’ (Mn-AOM) (Leu *et al*., 2020), *’Ca.* Methanoperedens nitroreducens’ Type Strain (electrode/nitrate/iron-AOM) (Zhang *et al*., 2023) and *’Ca.* Methanoperedens Vercelli’ (electrode-AOM) (Ouboter *et al*., 2024). The identified MHCs domains were subtracted from the protein sequence and aligned using MAFFT v7.525 (Katoh & Standley, 2013) via https://www.ebi.ac.uk/ with parameters --bl 62 -- op 1.53 --ep 0.123 --reorder --retree 2 --treeout --maxiterate 2 –amino. Finally, we constructed a phylogenetic tree using IQ-TREE v2.1.4 with function -s -st AA -m MFP -bb 1000 -nt AUTO, with a VT+R6 as best-fit model based on Bayesian Information Criterion (BIC). All presented trees were annotated using iTOL v5 (Letunic & Bork, 2021).

### Extrachromosomal elements (ECEs) search

We additionally screened for putative ECEs in our five ‘*Ca.* Methanoperedens’ MAGs. Among the ECEs investigated we included BORGs. For this, we first obtained a PFAM/InterPro database with dereplicated sequence representatives built from 40 different unique markers obtained from 17 different BORGs that appear to be associated with ‘*Ca.* Methanoperedens’ (Schoelmerich *et al*., 2024). Then we used DIAMOND v2.1.9.163 (Buchfink *et al*., 2015) to perform a BLASTp query search against our ‘*Ca.* Methanoperedens’ MAGs. For these annotations we applied an e-value cut-off of 10^-5^ and a minimum amino acid identity of 30%. We also searched for plasmids and viruses integrated into the ‘*Ca.* Methanoperedens’ MAGs described in this study by using VIBRANT (Kieft *et al*., 2020) and geNomad (Camargo *et al*., 2024) with default parameters. Initially, we identified 68 contigs as potential ECEs. To further investigate the nature of contigs, we classified them using specialized viral and phage annotation databases, including PHROGs (Terzian *et al*., 2021) pVOGs (Grazziotin *et al*., 2017), and VOGs (Bao *et al*., 2004). The e-value and identity cut-offs used for this annotation were 10^-5^ and 30%, respectively.

## Results

### Long-term incubations with different electron acceptors preserve and enrich ‘*Ca*. Methanoperedens’ as the key driver of AOM

The original carbon isotopic composition of the organic matter in the sediment used for the incubation experiment was -29.11±1.99 ‰ **δ**^13^C-TOC. As we used ^13^C-labeled methane to trace AOM in the slurry incubations with different electron acceptors (Supplementary Table 1), any change towards more positive **δ**^13^C-TOC values is caused by the production of new biomass from ^13^C-methane. Similarly, production of enriched **δ**^13^C-DIC indicates active AOM. Accordingly, all incubations showed more or less AOM activity, as indicated by the different enrichment in ^13^C. Most strikingly, **δ**^13^C-DIC in the incubation with added MnO_2_ was significantly enriched in ^13^C relative to the control or bottles with sulfate addition, whereas the δ¹³C value was less positive in the sample when both MnO_2_ and MoO_4_^2-^ were present (one order of magnitude lower than the control). Likewise, both TOC and TC had much more positive δ¹³C values in the solid phase of the MnO_2_-amended slurry compared to the incubation with sulfate addition or the unamended control. Notably, substantial amounts of sulfate were produced during the long-term incubation with MnO_2_, while sulfide remained close to the detection limit. By comparison, high concentrations of sulfide (∼ 0.5 mM) were detected in the slurry with sulfate addition at the end of incubation (Supplementary Table 1). By 16S rRNA gene sequencing at the end of the incubation period, we find that all incubations were dominated by bacteria, predominantly composed of Chloroflexi and Proteobacteria, including the phylum Aminecenantes (Supplementary Figure 1A and C). Within the archaeal fraction, Euryarchaeota dominated all incubations, particularly the manganese oxide incubation, followed by Woesearchaeota (Supplementary Figure 1A and B).

‘*Ca.* Methanoperedens’ was retained under all conditions and was most enriched (19.2% relative abundance) in the manganese oxide incubation (Supplementary Figure 1D). In contrast, incubation with both sulfate and molybdate had the lowest relative abundance (0.04%). In slurries amended with other electron acceptors, including the incubation with both manganese oxides and molybdate, the relative abundances of ‘*Ca.* Methanoperedens’ spp. were between 0.52% and 1.09%. All incubations selected for SRB closer to the *Syntrophaceae* rather than for the uncultured *Desulfobacterota* representative, previously described as *Desulfobulbaceae* (Supplementary Figure 1E).

### Co-occurrence of ‘Sed MAG Methanoperedens 1’ and SRB *Desulfobacterota* class *QYQD01* supports possible syntrophic interactions

To resolve the co-occurrence and potential interaction of ANMEs and SRB in Lake Cadagno, a metagenome analysis was conducted, and five ‘*Ca*. Methanoperedens’ MAGs were retrieved (Figure 2). Four of the *’Ca*. Methanoperedens’ MAGs were abundant in the original sediment while one MAG (labelled ‘MnO2 MAG Methanoperedens’, light blue) dominated the MnO_2_ incubation (Figure 2). The most abundant Methanoperedens MAG in the original sediment sample, ‘Sed MAG Methanoperedens 1’, contained a full-length 16S rRNA gene that showed 100% identity to one of the sequence variants recovered by Su *et al*. 2020 (Figure 1, ZOTU202 and Supplementary Figure 4 in original paper). ‘Sed MAG Methanoperedens 1’ sustained its dominance as the main anaerobic methane oxidizer for the unamended control and the sulfate incubation (Figure 2). The numbering of the ‘*Ca*. Methanoperedens’ MAGs is based on the metagenomic abundance of archaeal MAGs, from most to least abundant, in the co-assembled sediment metagenome (23 cm depth) (Supplementary Table 2). This resulted in ‘Sed MAG Methanoperedens 1, 2, 3 and 5’ (Figure 2), with ‘Sed MAG 4 c_Thermoplasmata’ being assigned number four.

**Figure 2.**
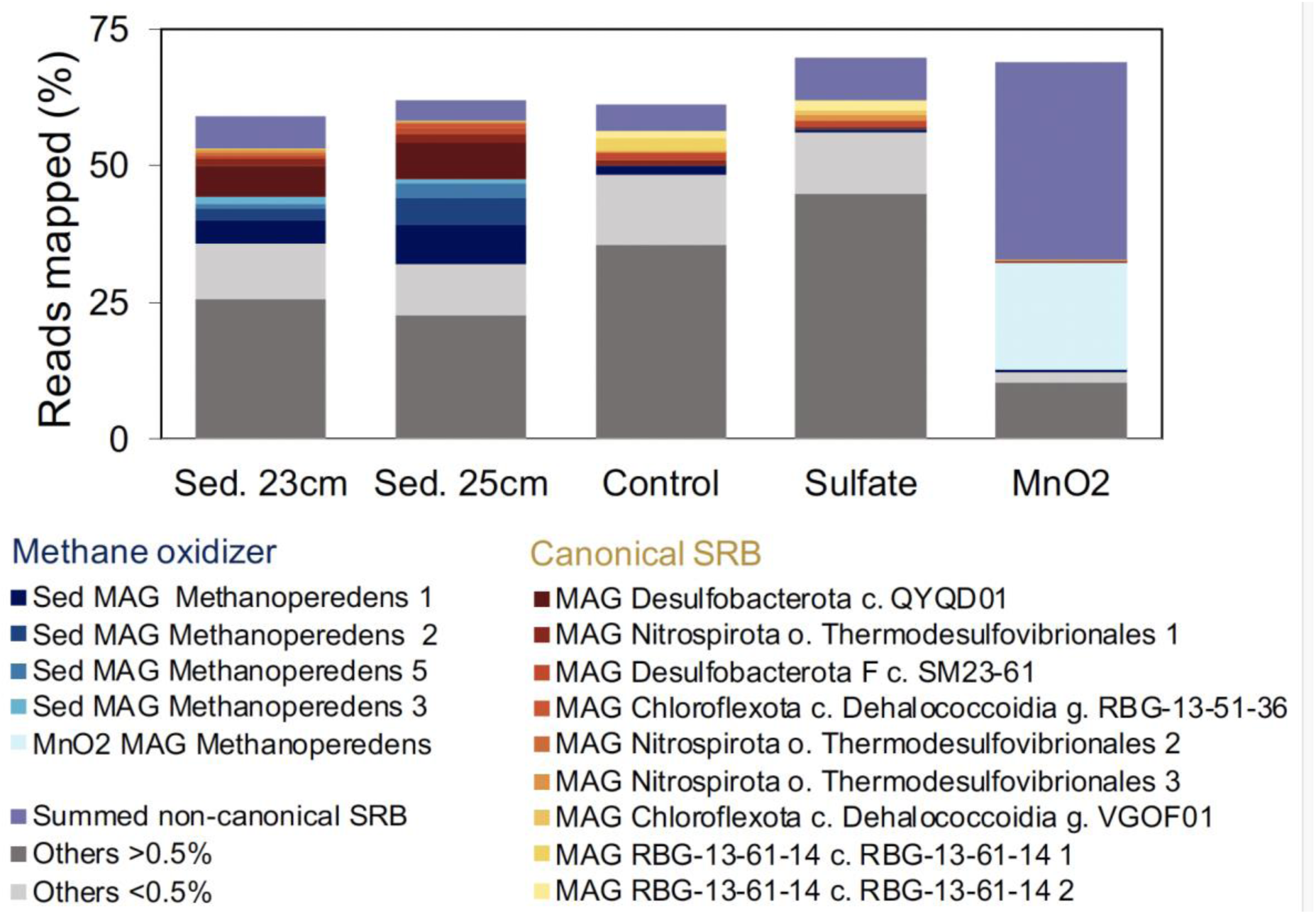
Percentage of mapped reads of top (above 0.5% for all metagenome sources) Metagenome Assembled Genomes (MAGs) analyzed across the two sediment depths (23 cm and 25 cm) and the three incubations: unamended control, sulfate, and manganese oxides. Subcategories include anaerobic methane oxidizers and canonical sulfate-reducing bacteria (SRB) (containing *dsrABCD*) and non-canonical SRB (containing *dsrABC*), as described in Diao *et al*. 2023. The ‘Others’ category is subdivided into all genomes that either exceeded 0.5% (dark grey) across all metagenome sources or fell below this threshold (light grey). The remaining reads constitute the unbinned fraction (not filling up to 100%).

The most prominent SRB MAG in the sediment belonged to the *Desulfobacterota* class *QYQD01*, with 5.6 to 6.7% of the reads in sediment 23 cm and 25 cm, respectively (Figure 2 and Supplementary Table 3). ‘MAG Desulfobacterota c. QYQD01’ showed 100% of 16S rRNA gene identity match with the most abundant uncultured Desulfobacterota ZOTU307, previously described as ‘uncultured *Desulfobulbaceae*’ (Su *et al*., 2020). The closest related microorganism to ‘MAG Desulfobacterota c. QYQD01’ in the NCBI database is the cultured *Desulfobacca acetoxidans* DSM11099, with an amino acid identity of 89.7% and 89% query cover. Consistent with the 16S rRNA gene analysis, the abundance of ‘MAG Desulfobacterota c. QYQD01’ decreased to 0.2%, 0.05%, and 0.03% in the unamended control, sulfate, and manganese oxide incubations, respectively (Figure 2, Supplementary Figure 1 and Supplementary Table 3). The unamended control and the sulfate treatment selected for canonical SRB from the order *Thermodesulfovibrionales*, as well as for the *Chloroflexota* phylum, genus *RBG-13-51-26,* or the phylum *RGB-13-61-14* (Supplementary Tables 3, 4 and 5).

### MnO_2_ amendments showed marked potential for Mn-AOM in sediment and strongly enriched for one ‘*Ca*. Methanoperedens’ sp. representative in the seed community

The most abundant MAG in the manganese oxide incubations, ‘MnO2 MAG Methanoperedens 1’ showed 99.7% 16S rRNA identity with one particular ‘*Ca.* Methanoperedens’ sp. ZOTU1150 (Su *et al*., 2020). When compared to MAGs ‘Sed MAG Methanoperedens 1,2,3 and 5’, the ‘MnO2 MAG Methanoperedens’ was rare in the original sediment, with around 0.007% relative abundance, and enriched to about 19.5 % in the incubations amended with manganese oxide (Supplementary Table 3 and 6). These incubations also favored the enrichment of non-canonical SRB from the Chloroflexota phylum and the order *Anaerolineales*. These organisms lack the *dsrD* subunit but include sulfide oxidoreductases (*sqr*) (classified as non-canonical SRB). The *Anaerolineales* contributed as much as 21% to the total metagenome (Supplementary Table 6).

### Recovered ‘*Ca.* Methanoperedens’ spp. show EET potential

To uncover the main metabolic traits and biogeography of the five Lake Cadagno ‘*Ca.* Methanoperedens’ MAGs, we generated an annotated genome tree (Figure 3).

**Figure 3.**
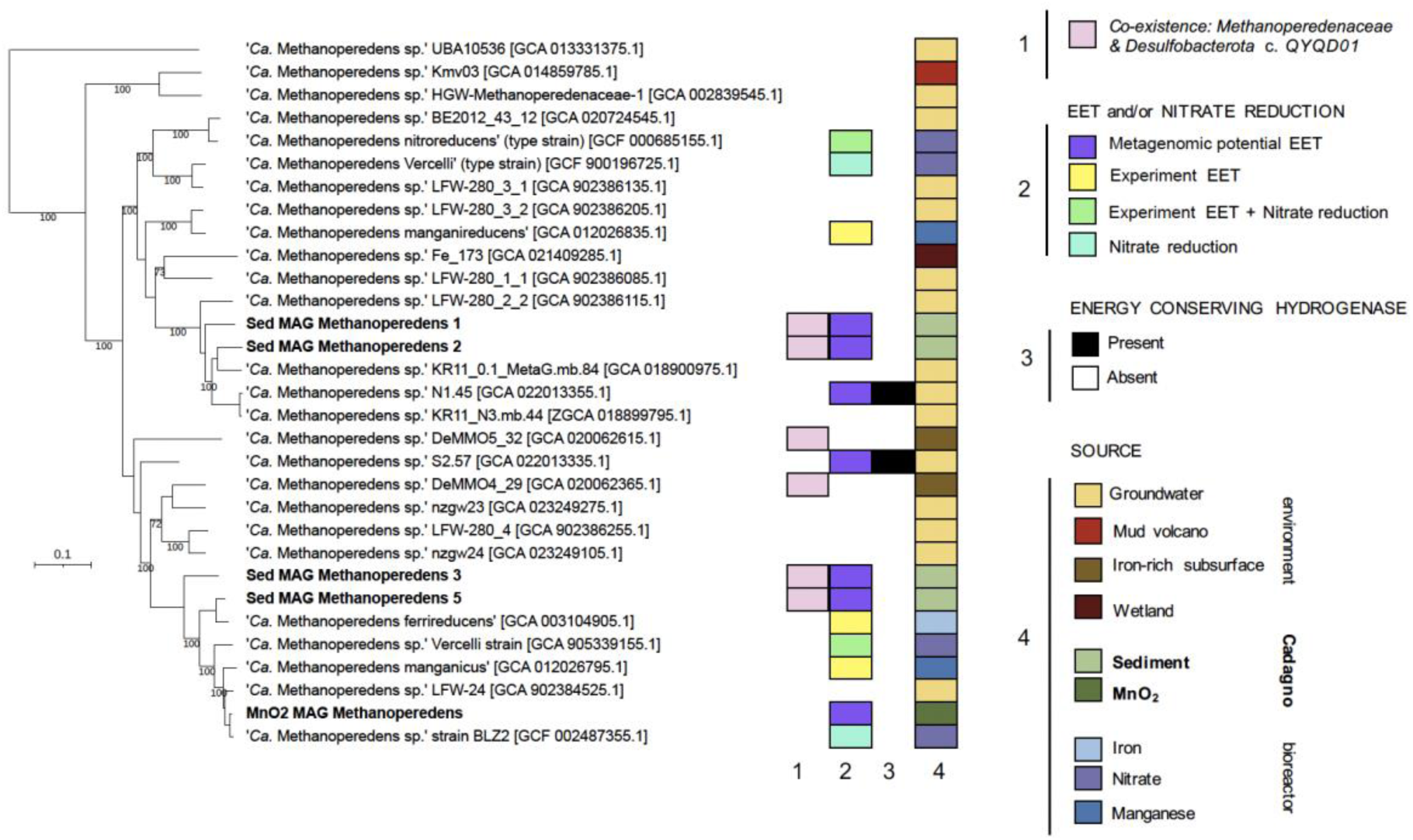
Phylogenomic tree of *Methanoperedeneacae* MAGs with Lake Cadagno representatives highlighted in bold. The tree has been generated using GTDB-Tk classification tools with multiple sequence alignments of 53 archaeal phylogenetic markers. Four MAGs that were not in the GTDB database at the time of analysis were additionally included: GCA_022013355.1 & GCA_022013335.1 (Bell *et al*., 2022) and GCA_020062615.1 & GCA_020062365.1 (Casar *et al*., 2021). Extracellular electron transfer (EET) and presence/absence of hydrogenases are only indicated for MAGs with shown or suspected EET. Branch lengths represent the average number of amino acid substitutions per site. Bootstrap values are shown for > 70% branching support.

‘Sed MAG Methanoperedens 1 and 2’ were most closely related to a groundwater ‘*Ca.* Methanoperedens’ spp. (Figure 3). Conversely, ‘Sed MAG Methanoperedens 3 and 5’ and ‘MnO2 MAG Methanoperedens’ were more similar to ‘*Ca.* Methanoperedens’ spp. from bioreactor enrichments amended with nitrate or metal oxides. Among them, ‘MnO2 MAG Methanoperedens’ and the nitrate-enriched ‘*Ca.* Methanoperedens BLZ2’ exhibit the greatest similarity, sharing an AAI of 92% (Figure 3 and Supplementary Figure 2). Although phylogenetically closely affiliated with nitrate-reducing species, none of the Lake Cadagno ‘*Ca.* Methanoperedens’ MAGs contained a nitrate reductase. We also screened for DNRA (dissimilatory nitrate reduction to ammonium) potential by searching for the functional genes for nitrite reductase (cytochrome c-552) (*nrfA*) and cytochrome *c* nitrite reductase small subunit (*nrfH*). In the sediment-associated *’Ca.* Methanoperedens’ MAGs, we did not detect any *nrfAH* genes. However, in the ‘MnO2 MAG Methanoperedens’, we identified two *nrfA* genes (the catalytic subunit), but no *nrfH* was found, even in the unbinned fraction.

All Lake Cadagno ‘*Ca.* Methanoperedens’ MAGs included the full reverse methanogenesis pathway for anaerobic oxidation of methane and lacked cytosolic or energy conserving hydrogenases (Ech) (Supplementary Table 7). Closer inspection of the Lake Cadagno ‘*Ca.* Methanoperedens’ MAGs revealed the potential for EET via MHC proteins or OmcZ-like subunits, possibly forming nanowires (Figure 3 and 4).

**Figure 4.**
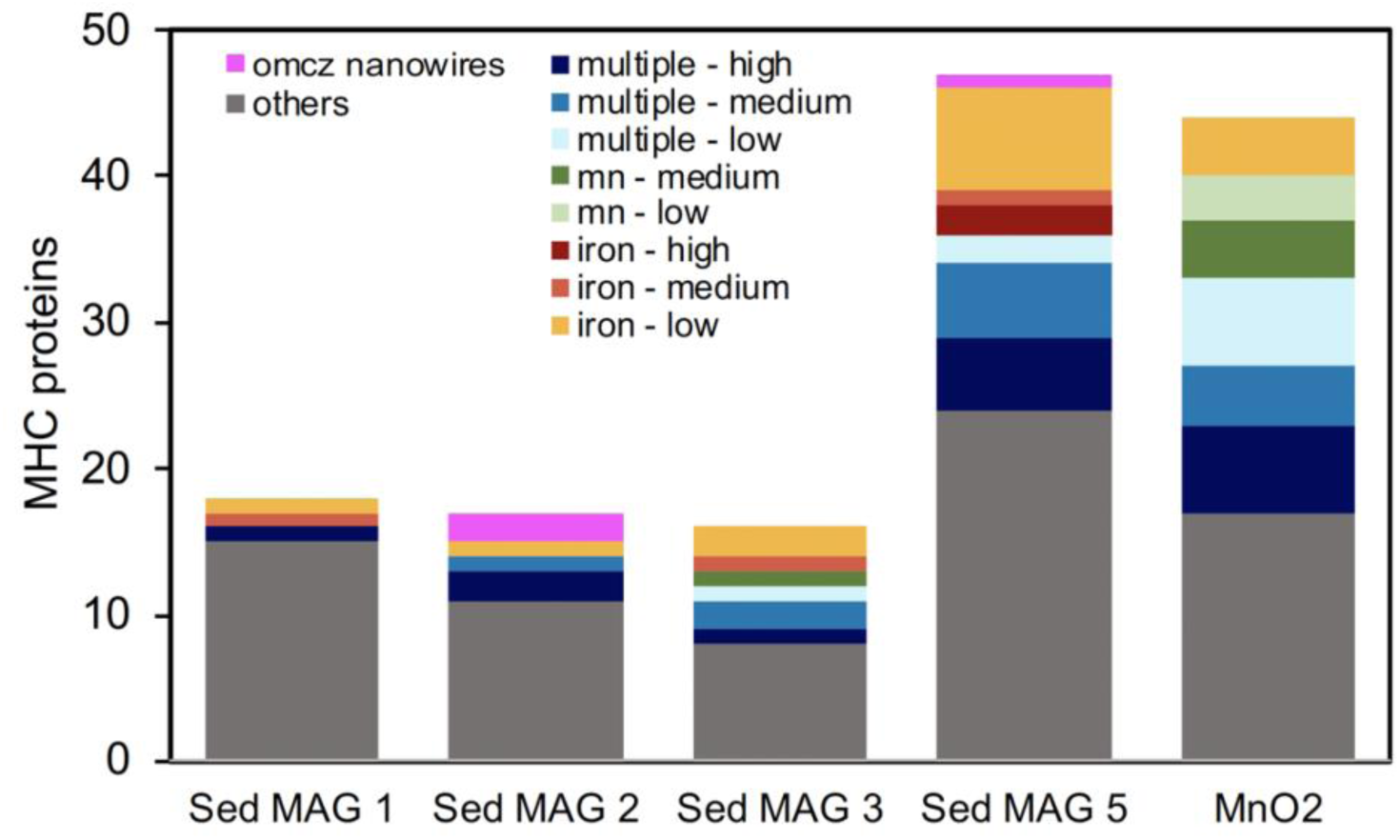
Multi-heme c-type cytochrome (MHC) protein counts in the Lake Cadagno ‘*Ca.* Methanoperedens sp.’ MAGs. Annotations are based on similarity (>70% to domain MHC) to reference ‘*Ca.* Methanoperedens’ enrichment species that showed MHC expression under different electron accepting conditions - multiple (including electrode), manganese or iron oxides – and expression levels. ‘High’ expression refers to the top 10% of the MHC transcripts, ‘medium’ to the following 40%, and ‘low’ to the remaining MHC transcript expression. OmcZ-like nanowire subunits are included as a separate category. The “Others” category includes MHC genes from this study that lack domain-homology-based expression links to reference ‘*Ca*. Methanoperedens’ MAGs. Comparing the similarity of MHC proteins among the Lake Cadagno ‘*Ca.* Methanoperedens’ spp. (Supplementary Figure 4) revealed that they shared between 9 and 16 homologous MHC proteins. For details see Supplementary Figure 4.

### Distinct multi-heme *c*-type cytochrome (MHC) and OmcZ-like nanowires assigned to Lake Cadagno ‘*Ca.* Methanoperedens’ spp

Intrigued by the high AOM activity in the manganese oxide incubation and the enrichment of ‘MnO2 MAG Methanoperedens’, we analyzed the metagenomes carefully for putative EET mechanisms. We investigated the amount, type, and similarity of MHC proteins plus their OmcZ-like subunits putatively forming conductive nanowire proteins (Figure 4, Supplementary Figure 3-4 and Supplementary Table 8). ‘MnO2 MAG Methanoperedens’ and ‘Sed MAG Methanoperedens 5’ showed almost twice the amount of MHC proteins compared to ‘Sed MAG Methanoperedens 1, 2 and 3’. ‘MnO2 MAG Methanoperedens’ and ‘Sed MAG Methanoperedens 5’ had 44 and 47 MHCs, respectively (Figure 4). Only ‘Sed MAG Methanoperedens 2 and 5’ encoded two and one OmcZ-like subunits possibly forming nanowire proteins, respectively. The other ‘MnO2 MAG Methanoperedens’ did not contain *omcZ* like genes. Instead, ‘MnO2 MAG Methanoperedens’ (contig_13_120) contained an Extracellular Cytochrome Nanowire (ECN) (Baquero *et al*., 2023) that shared 98% BLASTp identity with ‘*Ca.* Methanoperedens BLZ2’ ECN [WP_097300794.1], matching the high AAI (92%) of both MAGs (Supplementary Figure 2).

In Ouboter *et al*. 2024 and Leu *et al*. 2020, several gene clusters are described to be involved in EET by ‘*Ca.* Methanoperedens’. Therefore, we assessed which MHC genes of the Lake Cadagno ‘*Ca*. Methanoperedens’ MAGs resembled those clusters. We find that more than half of the MHC proteins in ‘MnO2 MAG Methanoperedens’ and in ‘Sed MAG Methanoperedens 3 and 5’ showed high homology with MHC proteins expressed in the reference ‘*Ca.* Methanoperedens’ spp. (Figure 4). For example, in ‘MnO2 MAG Methanoperedens’, 7 out of the 44 MHC proteins were related to those highly expressed in ‘*Ca.* Methanoperedens’ spp. grown with manganese oxides as the electron acceptor, while 4 were related to MHC proteins with low expression under iron-oxide-reducing conditions. In ‘Sed MAG Methanoperedens 5’, 9 out of the 49 MHC proteins corresponded to MHC proteins expressed in the iron-AOM ‘*Ca.* Methanoperedens’ enrichment, with the majority showing low expression levels, and a few matching those with medium or high expression (Figure 4). Furthermore, ‘Sed MAG Methanoperedens 5’ *omcZ*-like genes appeared to be clustering close to the highly expressed *omcZ*-like gene in ‘*Ca.* Methanoperedens ferrireducens’ performing Fe-AOM (Figure 4 and Supplementary Figure 3).

We also examined the presence of additional EET electroconductive structures. We identified type IV pilus assembly proteins in all Lake Cadagno ‘*Ca.* Methanoperedens’ MAGs. Furthermore, we investigated the formation mechanisms of extracellular polymeric substances (EPS), as these could facilitate the development of ANME/SRB consortia through the formation of cell aggregates and biofilms, as described by Murali *et al*. 2023. This search revealed the presence of cohesion subunit Scc3/SA, dockerin-like domain proteins and extracellular Contractile Injection Systems (eCIS) across all ‘*Ca*. Methanoperedens’ MAGs investigated, which either could help in the cellular adhesion or intercellular communication between organisms (Supplementary Table 9).

### Phylogenetic, biogeography and genomic analysis of the putative S-AOM partner Desulfobacterota class QYQD01

After the striking co-occurrence of the SRB *Desulfobacterota* class *QYQD01* with ‘*Ca.* Methanoperedens’ in Lake Cadagno sediments, we took a closer look at the *Desulfobacterota* class *QYQD01* MAG. We observed that the class *QYQD01* and the neighboring class *DTXEO1* form a distinct clade within the *Desulfobacterota* phylum, with the order *Desulfatiglandale*s representing the most closely related cultivated SRB (Figure 5A). We identified three additional *Desulfobacterota* class *QYQD01* genomes in the GTBD, two of which, DeMMO_14 (Deep Mine Microbial Observatory) and PowLak16 (Powell Lake), classified as the same species, and all three co-occurred with *Methanoperedenaceae* (Figure 5A and Supplementary Figure 6). Both the *Desulfobacterota* class *QYQD01* and *DTXEO1* genomes were recovered from diverse environments such as hydrothermal vents, sediments, meromictic lakes, or iron-rich subsurface waters. Notably, in the iron-rich subsurface water metagenome, Desulfobacterota MAG (DeMMO3_14) was found alongside a ‘*Ca.* Methanoperedens’ MAGs (DeMMO4_29 and DeMMO5_32; Figure 3).

**Figure 5.**
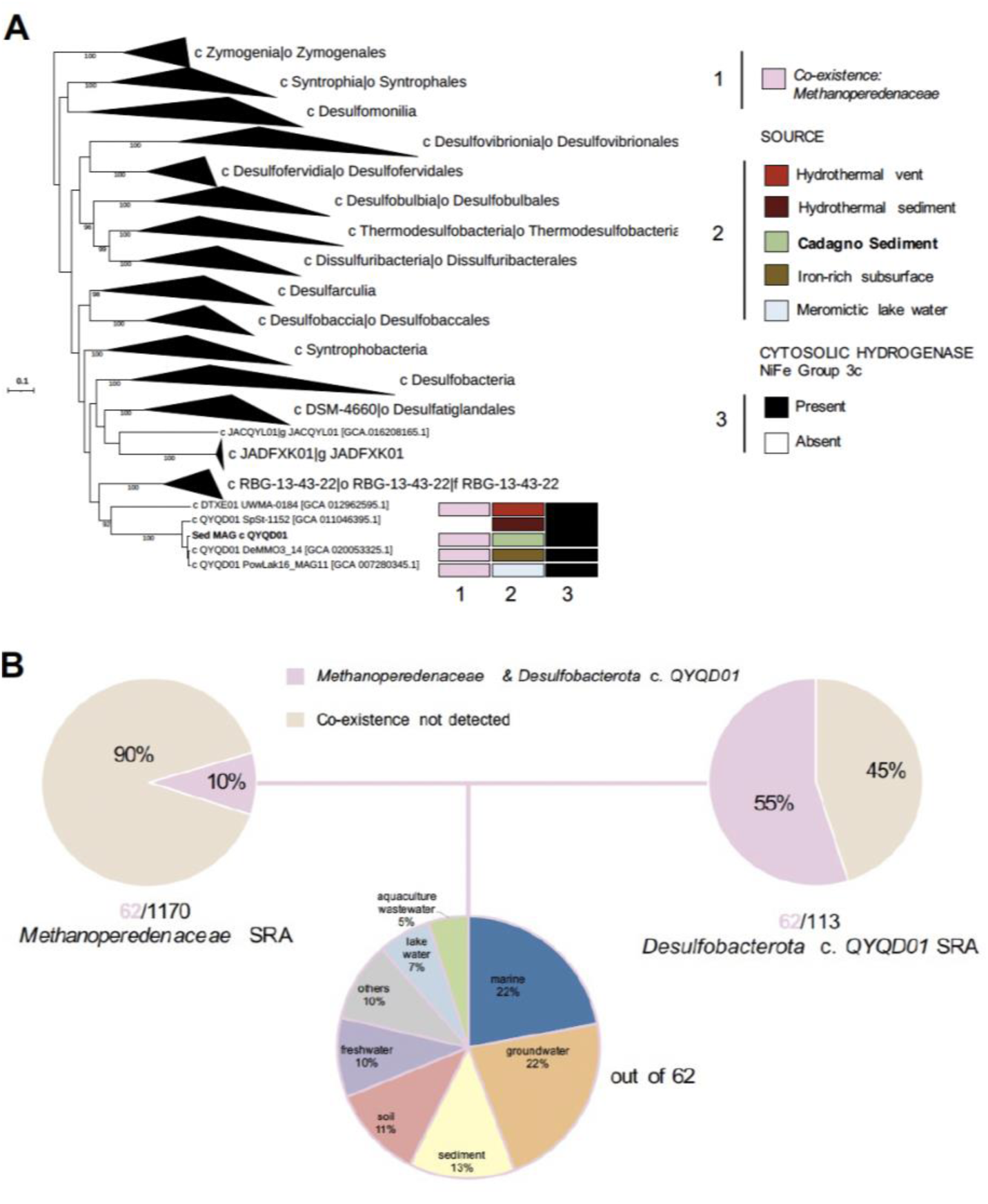
**A** Phylogenomic tree of the *Desulfobacterota* phylum based on Metagenome Assembled Genomes (MAG) with collapsed clades except for class QYQD01 and class DTXE01. The tree was generated via GTDB-Tk classification tools using multiple sequence alignments via concatenation of 120 bacterial phylogenetic markers. One MAG was included in addition to the genomes retrieved from the GTDB database: GCA_020053325.1 (Casar *et al*., 2021). Bootstrap values are shown for > 70% branching support. Branch length represents the average number of amino acid substitutions per site. **B** From top left to right, sequence read archive (SRA) hits for *Methanoperedeneaceae* family and *Desulfobacterota* class QYQD01. Colors indicate percentages of Sequencing Read Archive (SRA) entries where both groups were found in the same data set (pink) or not (brown). Bottom pie-chart displays SRA source from where both groups were found present.

Using the tool Sandpiper, we screened the sequence read archive (SRA) for metagenomic reads of *Desulfobacterota* class *QYQD01*, to assess whether this class has been observed alongside *Methanoperedenaceae* before (Supplementary Table 10 and 11). Our search resulted in 62 metagenomes where their co-existence was detected (Figure 5B). A quarter of the sequence SRA descriptions belonged to marine and groundwater systems, followed by sediment, soil, freshwater, as well as lake water, and aquaculture waste (Figure 5B). We determined the metagenomic coverage of reads from *Methanoperedenaceae* and *Desulfobacterota* class *QYQD01* in the 62 metagenomes in the ecosystems where they co-occurred (Figure 5B and Supplementary Figure 6A). The metagenomic coverage of *Methanoperedenaceae* to *Desulfobacterota* class *QYQD01* was much higher in groundwater systems (0 to 360) than in marine ecosystems (0 to 25) (Supplementary Figure 6A). We also found that *Methanoperedenaceae* co-existed with Desulfobacterota (class *QYQD01*) in about 70% of the groundwater samples from Finland’s Olkiluoto Island, and 21% of those from an arsenic-contaminated site in China (Supplementary Figure 6B). The Olkiluoto Island deep subsurface sample source was also shared by the *Methanoperedenaceae* MAGs presented in the above presented genome tree, labelled as KR11_0.1_MetaG.mb.84 & KR_11_N3.mb.44 (Mehrshad *et al*., 2021) and N1.45 and s2.57 (Bell *et al*., 2022) (Figure 3). The marine metagenome samples were dominated by mangrove, seagrass, estuary or oilfield sediments (50%), followed by the Deep Horizon Spill Sediment (21%) (Supplementary Figure 6B).

We additionally investigated the genetic features that could be indicative of a potential interdependency of the *Desulfobacterota* class *QYQD01* and ‘*Ca.* Methanoperedens’ spp. to sustain syntropy. The recovered *Desulfobacterota class QYQDO1* MAG contained the full respiratory sulfate reduction pathway that all ‘*Ca*. Methanoperedens’ spp. lacked. We found a total of 18 MHC encoded in the ‘MAG Desulfobacterota class QYDO1’ (Figure 5 and Supplementary Table 8). The largest MHC of ‘MAG Desulfobacterota class QYDO1’ was encoded within an operon comprising four adjacent sequences. These sequences encoded two MHC proteins - one with 12 heme binding motifs and another with 26 heme binding motifs - including a peptidyl-prolyl isomerase (or PPIase) sequence containing 5 heme binding motifs, and a small ORF of unknown function. The second gene cluster encoding large MHCs encoded two cytochromes with 11 and 12 heme-binding motifs, respectively, and an upstream Adenosine monophosphate (AMP) nucleoside-encoding protein plus two other cytochrome C assembly and biogenesis proteins (Supplementary Table 8).

Additionally, we also considered the syntrophic lifestyle of SRB associated with marine ANME, and included putative extracellular polysaccharides and protein complexes that could aid in the interaction with ‘*Ca*. Methanoperedens’ as described in Murali *et al*. 2023 (Supplementary Table 12). Consistent with our observations on the aggregate formation mechanisms of ‘*Ca*. Methanoperedens’ (Supplementary Table 9), the presence of certain marker genes in the ‘MAG Desulfobacterota class QYQD01’ supports ANME/SRB interaction and communication (Supplementary Table 12). These structures include type IV pili (EET mechanism), the type VI secretion system (for intercellular communication), and potential adhesins for cellular adhesion, such as the trimeric autotransporter adhesin YadA-like head domain (Supplementary Table 12).

We also analyzed some hallmark genomic traits that could suggest a free-living lifestyle of *Desulfobacterota* class QYQD01 as a putative autotrophic SRB. The ‘MAG Desulfobacterota c. QYQD01’ included the potential for carbon fixation via the Wood–Ljungdahl pathway. We also recovered non-energy conserving cytosolic hydrogenases (NiFe group 3c) for our study’s MAG and the three other *Desulfobacterota* class *QYQD01* MAGs included in the genome tree (Figure 5A). For our ‘MAG Desulfobacterota c. QYQD01’, the accessory hydrogenase subunits (*hyd*) appeared together in a separate region of the genome forming a different operon that was not adjacent to the large and small NiFe group 3c subunits.

Other observation included the lack of phosphate acetyltransferase and acetate kinase for conversion of acetyl-CoA to acetate and nitrate or nitrite reductase (Supplementary Table 7).

### Presence of Mobile Genetic Elements (MGEs) in Lake Cadagno’s ‘*Ca*. Methanoperedens’

To assess the metabolic flexibility and putative horizontal gene transfer (HGT) in the Lake Cadagno ‘*Ca*. Methanoperedens’ MAGs, we examined the presence of ECEs. In this regard, we addressed the presence of BORGs, which have been described as novel giant ECEs, that are not classifiable as virus or plasmid. They have been shown to be associated with ‘*Ca*. Methanoperedens’ and have propensity to assimilate genes that harbor potential for key metabolic pathways such as anaerobic methane oxidation, extracellular electron transfer, or stress resistance (Al-Shayeb *et al*., 2022, Schoelmerich *et al*., 2024). More recently, 40 unique putative BORGs markers have been described (Schoelmerich *et al*., 2024), which we employed against our ‘*Ca*. Methanoperedens’ MAGs. Our search resulted in 16 out of 30 family domain putative BORG marker proteins (Supplementary Table 13 and 14), suggesting the absence of previously described BORG ECEs in our samples.

We also explored additional mobile genetic elements (MGEs) belonging to the ‘*Ca*. Methanoperedens’ MAGs (Supplementary Tables 15-17). Our search identified 64 contigs as potential MGEs (Supplementary Table 18). To assess whether any of these were viral, we screened for specific structural proteins (e.g., capsids) but found no positive hits (Supplementary Tables 19-21). We then looked for alternative signature proteins, including integrases and transposases, detecting seven and six hits, respectively (Supplementary Tables 18-21). Finally, in those integrases and transposases harboring contigs, we checked for the presence of terminal repeats, an indicator of MGE completeness, and identified two instances in ‘Sed MAG Methanoperedens 1’ (contig_3571) and ‘Sed MAG Methanoperedens 2’ (contig_21328) (Supplementary Table 18).

We also evaluated the possible role of MGE in HGT of key genes associated with the adaptation of SRB into a syntrophic partnership with ANME, as proposed by Murali *et al*. 2023. However, proteins encoding structures responsible for aggregate formation between ANME and SRB (Supplementary Table 9) were not identified in any of the 64 potential MGEs investigated (Supplementary Table 18). This suggests that MGEs do not influence the establishment of this partnership.

## Discussion

This study deepens our knowledge of the poorly explored potential syntrophic interaction between ‘*Ca.* Methanoperedens’ spp. and the *Desulfobacterota* class *QYQD01* in Lake Cadagno sediments (Figure 1-2 and Supplementary Figure 1). After screening of the SRA, we also described the potentially widespread co-existence of *Desulfobacterota* class *QYQD01* and ‘*Ca.* Methanoperedens’ spp. (Figure 3 and 5) across many groundwater and marine ecosystems (Figure 5B and Supplementary Figure 6).

Long-term microcosm incubations under different electron acceptor conditions resulted in the enrichment of ‘MnO2 MAG Methanoperedens’, which constituted up to about 19.5% (Figure 3) of the microbial community, which aligns well with the almost ∼20% abundance of one of the ‘*Ca*. Methanoperedens’ ASVs observed (ZOTU1150) by Su *et al*. 2020. Molybdate addition to the manganese oxide incubation resulted in strong AOM inhibition and disappearance of the highly enriched ‘*Ca*. Methanoperedens’ spp. (Supplementary Table 1 and Supplementary Figure 1). This observation, coupled to the increase in non-canonical *sqr* harboring SRB from the class *Anaerolineales* (including *sqr*) (Figure 1 and Supplementary Table 6) for the same enrichment, is suggestive of cryptic sulfur cycling. Sulfate can be produced from reoxidation of sulfide or sulfide minerals present in the sediment with MnO_2_ (Supplementary Table 1) (Yao & Millero, 1996) allowing indirectly for sulfate reduction to occur. All incubations showed a loss of putatively syntrophic ‘MAG *Desulfobacterota* class *QYQD01’* (Figure 2). This is especially surprising for the sulfate enrichment, but the build-up of sulfide (488.6 µM) could explain the general sulfide-toxicity derived inhibition observed for other ANMEs (Supplementary Table 1) (Dalcin Martins *et al*., 2022).

Recently, Group III Dsr-LP sulfite reductases have been linked to sulfide toxicity in a ‘*Ca*. Methanoperedens BLZ2’ enriched culture (Echeveste Medrano *et al*., 2024b) and to sulfite detoxification in the model methanogen *Methanococcus maripaludis* (Day Leslie *et al*., 2024). In the current study, we recovered one Group III Dsr-LP protein in ‘Sed MAG Methanoperedens 1’ and ‘Sed MAG Methanoperedens 5’ and two for ‘Sed MAG Methanoperedens 2’ and the enriched ‘MnO2 MAG Methanoperedens’ representative. Group III Dsr-LP have been only described in some methanogens and ‘*Ca*. Methanoperedens’ spp., but not in marine ANME (Yu *et al*., 2018). Consistent with Echeveste Medrano *et al*. 2024b, we hypothesize that, given the absence of both nitrate and nitrite reductases (*nrfAH*) in the sediment ‘*Ca*. Methanoperedens’ MAGs and the high sulfate-to-nitrate availability, the putative role of Group III Dsr-LP sulfite reductases in the recovered MAGs is most likely linked to sulfite detoxification.

Our data suggest that ‘Sed MAG Methanoperedens 1’ and ‘Sed MAG Methanoperedens 2’ are the most plausible candidates to engage in a syntrophic interaction with *Desulfobacterota* class *QYQD01*. They clustered closer to environmental groundwater ‘*Ca.* Methanoperedens’ spp., which have more sulfate available, and some had also been found to co-occur with the *Desulfobacterota* class *QYQD01* (Figure 3). Conversely, ‘MnO2 MAG Methanoperedens’, ‘Sed MAG Methanoperedens 3’ and ‘Sed MAG Methanoperedens 5’, appeared to be more closely related to metal-reducing ‘*Ca.* Methanoperedens’ species (Figure 3). The putative MHC-enabled EET mechanisms of ‘MnO2 MAG Methanoperedens’ and ‘MAG Methanoperedens 5’ resemble most those of ‘*Ca.* Methanoperedens’ spp. enriched in cultures amended with metal (manganese and iron) oxides or other electron acceptors (electrode and nitrate) enrichments (Figure 3) (Cai *et al*., 2018, Leu *et al*., 2020, Zhang *et al*., 2023, Ouboter *et al*., 2024). On the contrary, the predicted syntrophic ‘Sed MAG Methanoperedens 1’ and ‘Sed MAG Methanoperedens 2’ harbor less homologous MHC proteins to that of known species, potentially indicating novel functionality (Figure 4 and Supplementary Figure 3). These two candidate MAGs were also more closely related to on the ones observed in the Olkiluoto Island deep subsurface (Figure 3).

Our co-occurrence *Desulfobacterota* class *QYQD01* and *Methanoperedenaceae* biogeography study revealed a high correlation of *Methanoperedenaceae* to *Desulfobacterota* class *QYQD01* in groundwater systems (Figure 5B and Supplementary Figure 6). This putative syntropy could represent a survival strategy for *Methanoperedenaceae* to dispose of electrons from methane oxidation when the environmental metal oxide pool gets depleted. One of the iron-rich subsurface metagenomic studies included genomes of both *Desulfobacterota* class *QYQD01* and two ‘*Ca.* Methanoperedens’ spp. labelled as DeMMO (Figure 4A and 5A) (Casar *et al*., 2021). They observed several genes encoding for proteins involved in iron cycling in six different fracture fluids with varying chemistry. Here, one clear difference between sites was the high sulfate concentration, ranging from 0.88 mM to 42.79 mM. The *Methanoperedenaceae* appeared to contribute to iron reduction the most in site D6, where sulfate was the most available from all sites. In Casar *et al*. 2021, SRB genomes from the ‘*Desulfobacterales*’ taxa were also suggested to contribute with 2-4% to the relative metagenomic iron reduction. The second *Desulfobacterota* class *QYQD01* genome (PowLake16_MAG 11) (Figure 5A) was sampled from the meromictic Powell Lake resembling Lake Cadagno as well as co-occurred with ‘*Ca*. Methanoperedens’ species (Supplementary Figure 6A, indicated with an arrow) (Haas *et al*., 2019).

We further explored ‘MAG Desulfobacterota class QYQD01’ for genomic features that could be indicative of a syntrophic lifestyle with ‘*Ca.* Methanoperedens’ spp. Our observations resulted in high congruence with genomic traits described for syntrophic SRB partners of marine ANME (Skennerton *et al*., 2017, Murali *et al*., 2023), including: the putative loss of Ech hydrogenase, type IV pili formation and type VI secretion system, presence of adhesins and the conservation of large MHCs for DIET. In this regard, the largest MHC in our ‘MAG Desulfobacterota class QYQD01’ was contained in a four-sequence operon structure that resembled the one described for SEEP-SRB 1 marine SRB as well as homologous organisms presented in the same study (Sed_Bac_MAG_1_p_Desulfobacterota_c_QYQD01_contig_10778 ORF, 1 to 4) (Supplementary Table 7 and 8). The analyzed SEEP-SRB MHC operon structure included one 26 and another 16 heme *c*-type cytochromes followed upstream by a PPIase domain and downstream by a beta propeller fold protein (Skennerton *et al*., 2017). Our ‘MAG Desulfobacterota class QYQD01’ also included a 26 and 12 (not 16) heme *c*-type cytochrome with a PPIase domain but since the contig in our study/MAG was broken downstream, we were not able to conclude whether this operon also included a gene encoding a beta-propeller fold protein. Compared to the genomic signature traits of marine SRB associated with ANME, our SRB lacked Ech hydrogenases. This observation contrasts with the NiFe group 1b hydrogenase reported for the Olkiluoto Island Deep subsurface in Finland, where sulfate-AOM was proposed via *Desulfobacterales* family *ETH-SRB1* and ‘*Ca.* Methanoperedens’ spp. N1.45 and S2.57 (Figure 3). For the here described putative syntrophic SRBs, we recovered cytosolic NiFe group 3c hydrogenases.

Murali *et al*. 2023 assembled widespread syntrophic marine SRB clades that partner with ANME to perform S-AOM and identified traits that might be suggestive of an adaptation to a syntrophic lifestyle. Here, additional features indicative of syntrophy were highlighted, such as the ability for biofilm formation, intercellular communication, or for some Seep-SRB1a, a nutritional dependency on ANME based on the lack of a cobalamin synthesis pathway. Our study’s *Desulfobacterota* class *QYQD01* clustered closely to marine syntrophic group Seep-SRB1g in the presented *Desulfobacterota* genome tree (Skennerton *et al*., 2017, Murali *et al*., 2023). For this comparative genomics study, *Desulfobacterales* family ETH-SRB1 (refer to as Seep-SRB1a sp.1) was also considered as a syntrophic SRB, aligning with the presented sulfate-AOM in co-abundance with ‘*Ca*. Methanoperedens’s spp. N1.45 and S2.57 in Olkiluoto Island deep subsurface in Finland (Bell *et al*., 2022, Murali *et al*., 2023).

Our investigation into ECEs in ‘*Ca*. Methanoperedens’ produced inconclusive results regarding the presence of BORGs. Specifically, the markers used to identify BORGs (Schoelmerich *et al*., 2024) did not provide clear evidence of their presence in the recovered genomes. Only 17 out of the 30 BORG family protein markers were detected (Supplementary Tables 13 and 14) so the presence of previously identified BORGs seems unlikely. Whether those 17 BORG family marker proteins belong to as of yet unidentified BORGs needs to be further investigated. Additionally, only two non-viral MGE belonging to ‘Sed MAG Methanoperedens 1’ and ‘Sed MAG Methanoperedens 2’ were conclusively identified (Supplementary Table 18).

To conclude, we describe an ubiquitous co-abundance of ‘*Ca.* Methanoperedens’ spp. and *Desulfobacterota* class *QYQD01* to sustain sulfate-AOM in sulfate-rich freshwater systems, mainly present in groundwater and marine systems. We suggest putative MHC-proteins of ‘*Ca.* Methanoperedens’ that could engage via EET in a syntrophic interaction and present metabolic adaptations and phylogenomic placement of *Desulfobacterota* indicative of a syntrophic lifestyle with ANME archaea. Future efforts should focus on S-AOM-targeted enrichments of ‘*Ca*. Methanoperedens’ and syntrophic SRB in bioreactors inoculated with sediment from Lake Cadagno. Metatranscriptomic and metaproteomic studies could be conducted on the sediment to determine which ‘*Ca*. Methanoperedens’ and SRB species are most active, identify the preferred MHC proteins, and investigate whether Group III Dsr-LP plays a role in sulfide-derived sulfite detoxification. Furthermore, analyzing the mineralogy of Lake Cadagno could provide insights into other available metal electron acceptors for ‘*Ca*. Methanoperedens’ spp. *in situ*.

## Acknowledgements

We thank Andy Leu for the bioinformatic data support and discussion. We acknowledge Thomas Kuhn for carbon isotopic analysis. Guangyi Su acknowledges a personal stipend from the China Scholarship Council (CSC). Additional funding came from the Department of Environmental Sciences of the University Basel. This study was supported by the Dutch Research Council (NWO) through the Gravitation Grant SIAM [Grant number 024.002.002] and an NWO-VIDI Talent grant [Grant number VI.Vidi.223.012]. It was furthermore supported by the ERC Synergy Grant MARIX [Grant number 854088].

## Data availability

Raw sequence data for 16S rRNA amplicon sequencing are made available at NCBI under the BioProject ID PRJNA497531 with the accession numbers SRR15689035-SRR15689041. Raw metagenomic data and MAGs recovered from the sediment and manganese oxide incubation co-assemblies have been uploaded to the European Nucleotide Archive with project number PRJEB81702. Supplementary Tables for this manuscript have been deposited in the Zenodo repository: https://zenodo.org/records/14055789

## Supplementary Data

**Supplementary Tables 2-21 (as a single spreadsheet)**

**Supplementary Table 1.** Chemical characterization of long-term incubated slurry samples in both aqueous and solid phases. Homogenized Lake Cadagno sediment of 19-24 cm was used as the inoculum for the slurry incubations, which were amended with ^13^C-labeled methane and different electron acceptors.

| Incubation samples |  | Unamended Control | Nitrate* 0.5 mM | FeO<sub>x</sub> 10 mM | MnO<sub>2</sub> 10 mM | MnO<sub>2</sub> + MoO<sub>4</sub><sup>2-</sup> 10 mM + 20 mM | Sulfate 4.8 mM | Sulfate + MoO<sub>4</sub><sup>2-</sup> 4.8 mM + 20 mM |
| --- | --- | --- | --- | --- | --- | --- | --- | --- |
| Aqueous phase | Sulfate (μM) | 0.6 | 0.8 | 0.2 | 272 | 320 | 105 | 320 |
|  | H<sub>2</sub>S (μM) | 42.2 | 49.9 | 1.1 | 0.8 | 0.8 | 488.6 | 38.2 |
|  | Fe<sup>2+</sup> (μM) | n.d. | n.d. | 172.1 | n.d. | n.d. | n.d. | n.d. |
|  | Mn<sup>2+</sup> (μM) | n.d. | n.d. | 7.3 | 337.8 | 420 | n.d. | n.d. |
|  | DIC (mM) | 2.67 | 3.2 | 5.71 | 0.92 | 0.66 | 5.70 | 1.96 |
|  | δ<sup>13</sup>C-DIC (‰) | 6651 | 6113 | 4215 | 16963 | 446 | 5849 | 3560 |
| Solid phase | TOC (%) | 2.24 | 2.55 | 3.03 | 2.68 | 2.21 | 2.98 | 2.89 |
|  | δ<sup>13</sup>C-TOC | -14.6 | -14.4 | -14.0 | 704.3 | -22.8 | -11.6 | -26.48 |
|  | TC (%) | 2.24 | 2.55 | 3.03 | 2.68 | 2.21 | 2.98 | 2.89 |
|  | δ<sup>13</sup>C-TC | -14.3 | -13.8 | -12.1 | 3442.4 | 49.5 | -11.1 | -26.3 |

n. d. = not detected.
\* Nitrate was only amended once at the beginning of incubation. Neither nitrate nor nitrite could be detected at the end of the incubation. Both compounds were also absent in all other incubation and are hence not reported in this table.

**Supplementary Figure 1.**
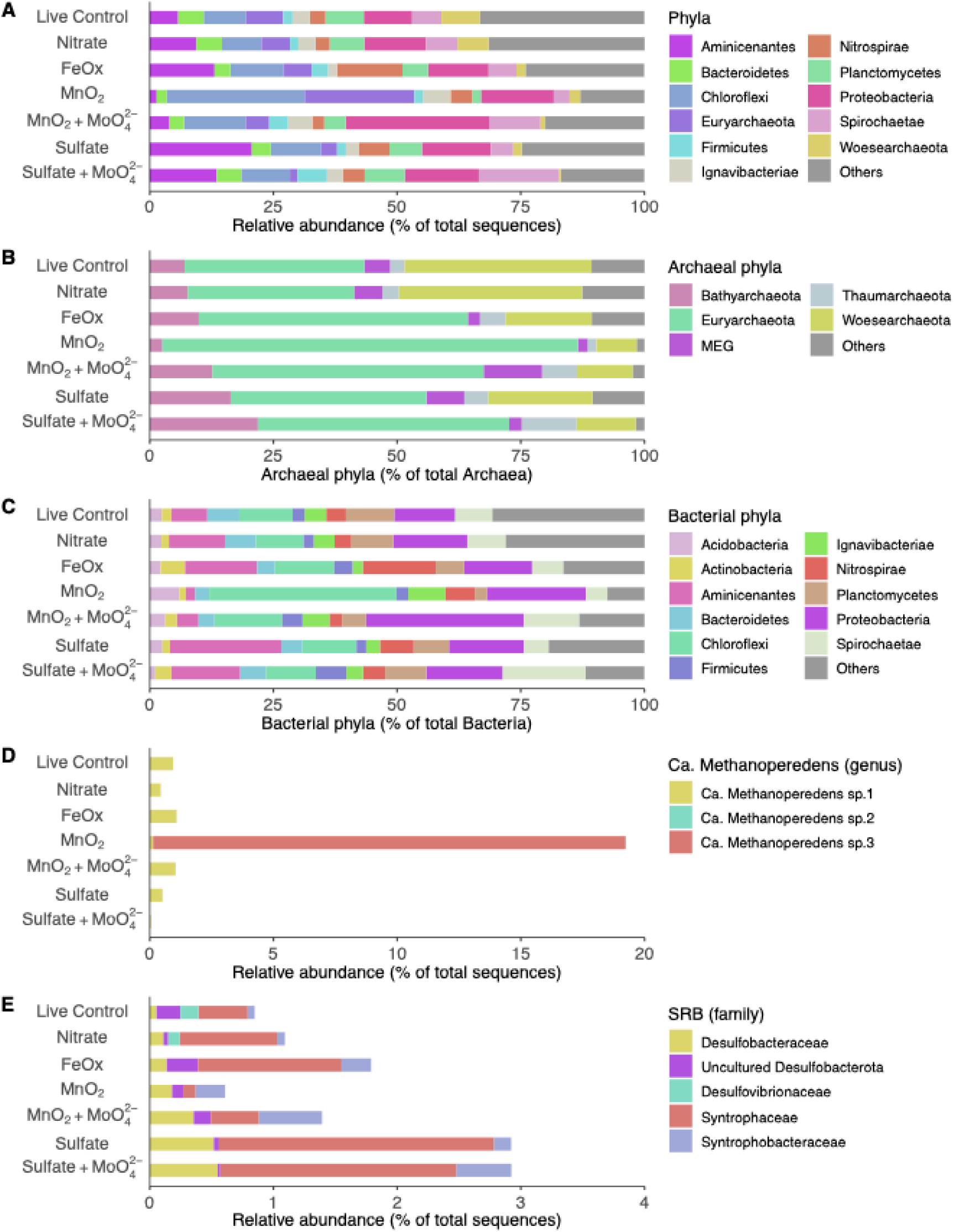
Microbial community composition at the end of the long-term incubation experiment, based on relative abundances of 16S rRNA gene amplicons. **A** Major phyla, **B** Archaea, **C** Bacteria**, D** ‘*Ca.* Methanoperedens’ genus Amplicon Sequencing Variants (ASVs), and **E** Potential sulfate-reducing bacteria (SRB).

**Supplementary Figure 2.**
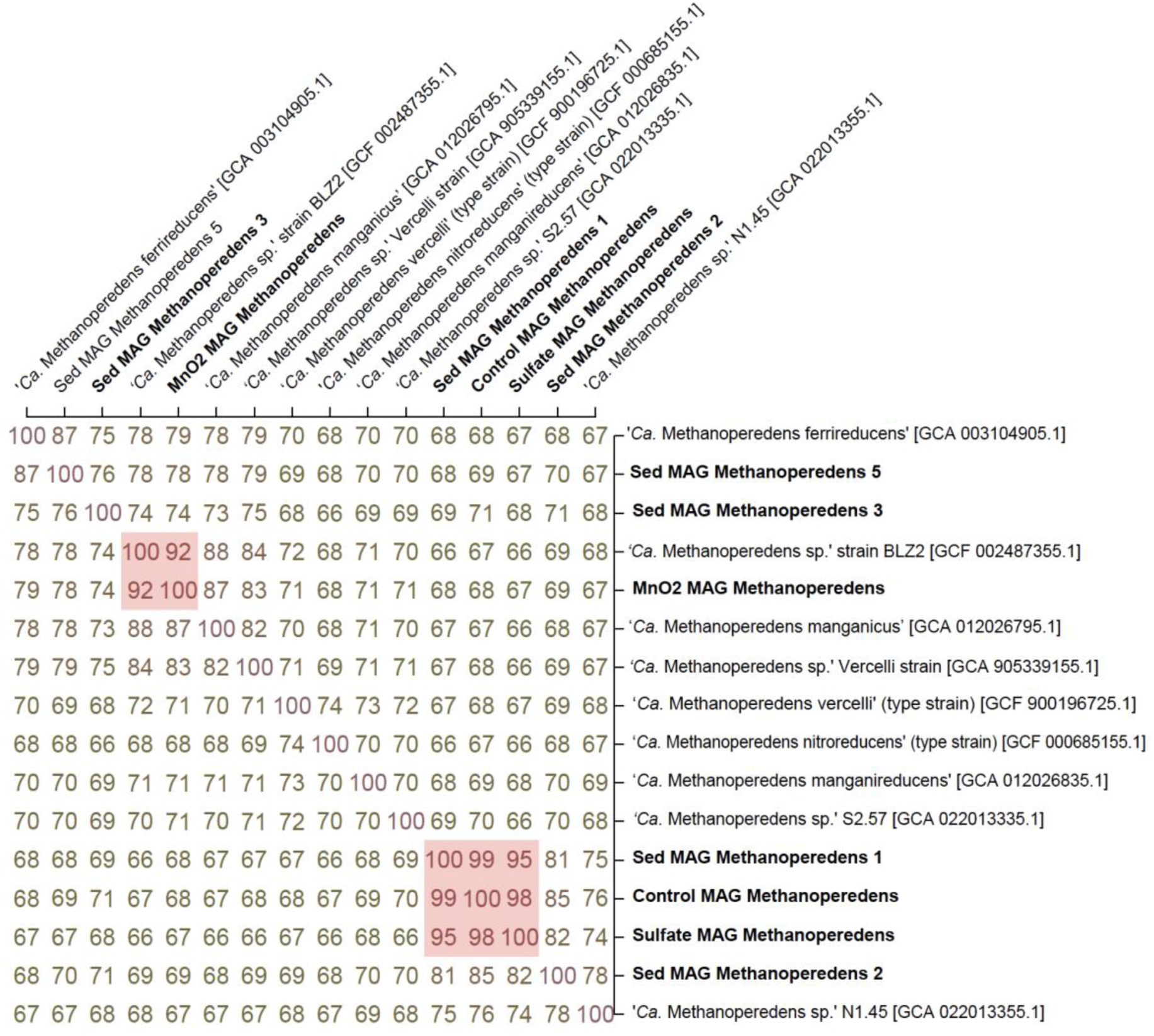
Average amino acid identity (AAI) matrix built with the current study’s ‘*Ca.* Methanoperedens’ spp. MAGs as well as MAGs belonging to reference bioreactor ‘*Ca.* Methanoperedens’ spp. enrichments. MAGs that are considered same species are highlighted in red (≥90% AAI) (Konstantinidis *et al*., 2022). MAGs are labelled with the given NCBI name id (biosample reference) and identifier in squared brackets.

**Supplementary Figure 3.**
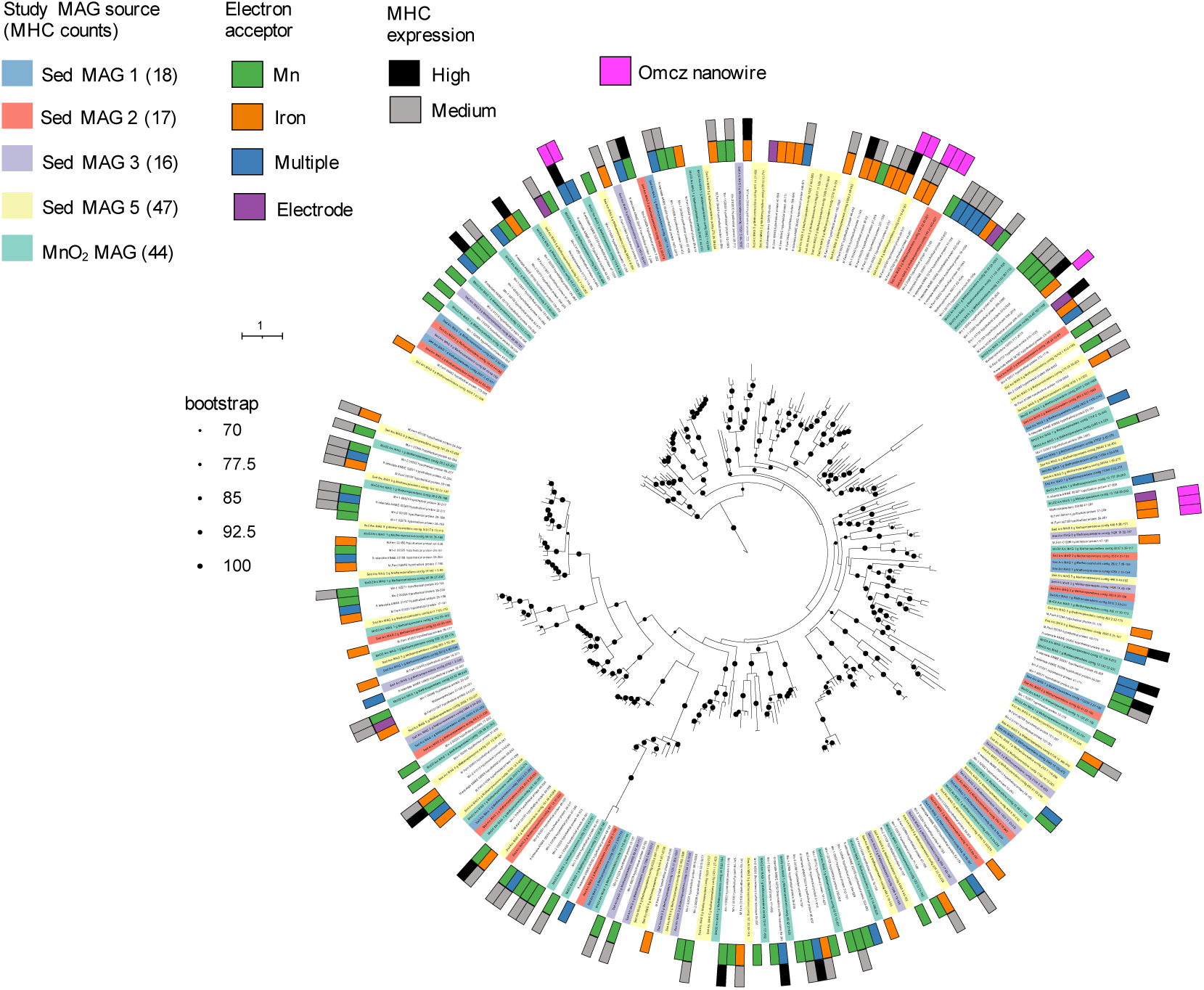
Multi-heme *c*-type cytochromes (MHC) family domain tree including current study’s ‘*Ca.* Methanoperedens’ MAGs and the reference ‘*Ca.* Methanoperedens’ bioreactor enrichments: *’Ca.* Methanoperedens ferrireducens’ (Fe-AOM) (ferri), *’Ca.* Methanoperedens manganicus (Mn-1)’ and *’Ca.* Methanoperedens manganireducens (Mn-2)’ (Mn-AOM), *’Ca.* Methanoperedens nitroreducens’ Type Strain (retentate) (electrode/nitrate/iron-AOM), *’Ca.* Methanoperedens Vercelli’ (electrode-AOM) (Methanoperedens). In the legend, from left to right: MAGs labelled based on current study origin (with total MHC domain counts in parenthesis), reference MAGs colored based on electron acceptor amendment (manganese oxides, iron oxides, multiple or electrode) as well as MHC expression (either high or medium) of the reference MAG proteins and proteins classified as nanowires (OmcZ). Branch lengths indicate the average number of amino acid substitutions per site. Black dots at nodes indicate robust branching with bootstrap values > 70%.

**Supplementary Figure 4.**
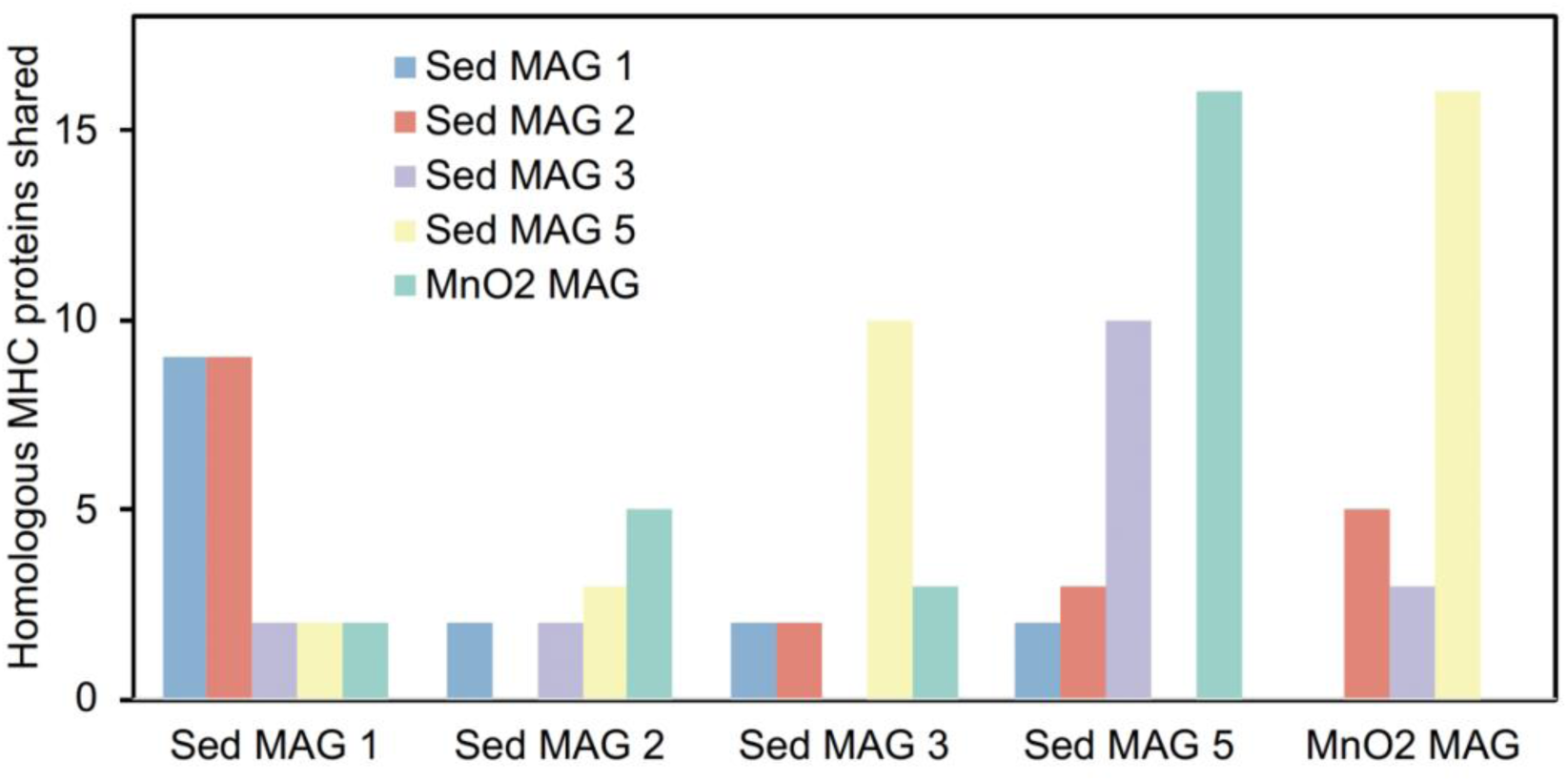
Multi-heme *c*-type cytochromes (MHC) family domain similarity among the Lake Cadagno ‘*Ca.* Methanoperedens’ MAGs. We considered homologous protein when the amino acid similarity was >70% of the MHC domain.

**Supplementary Figure 5.**
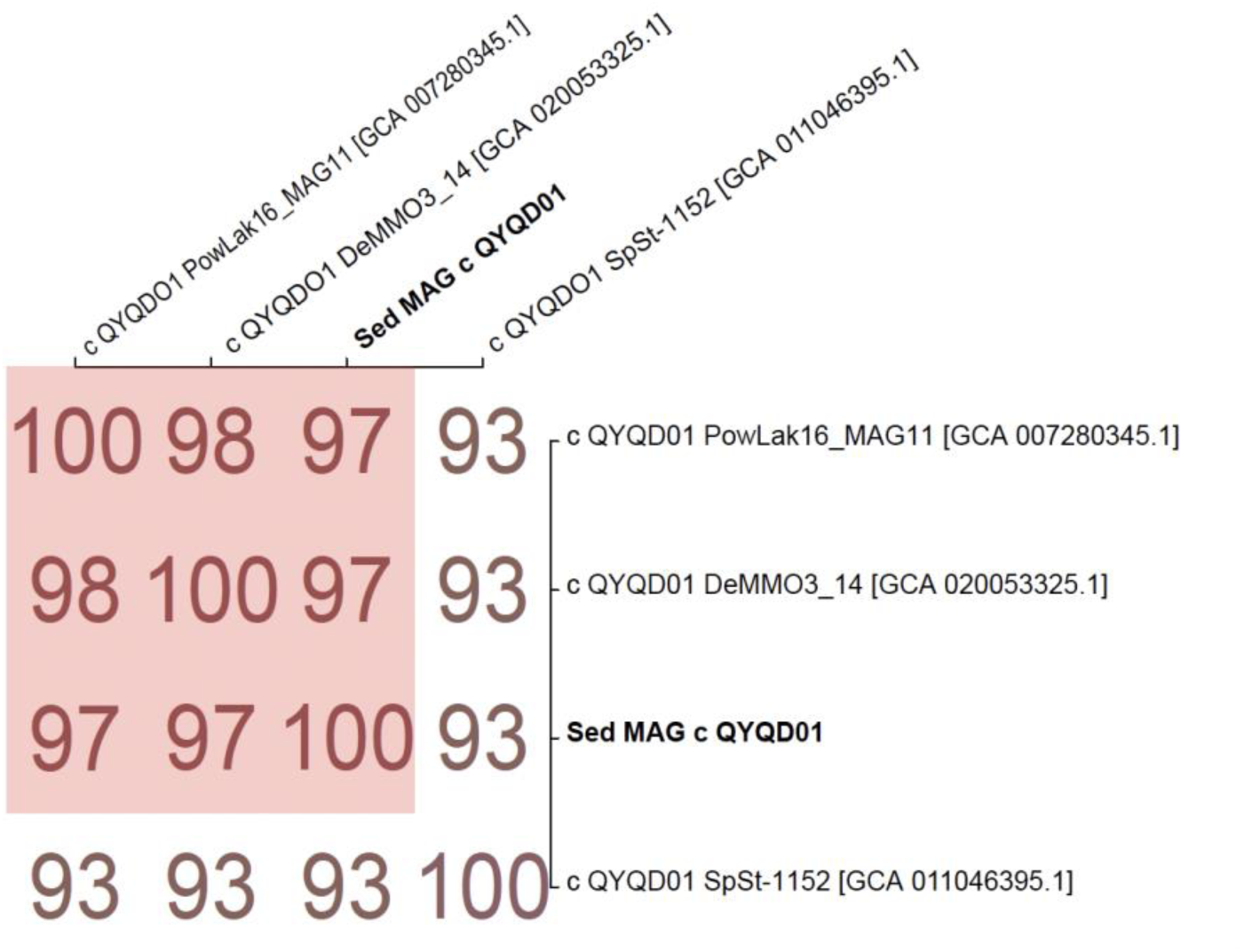
Average nucleotide identity (ANI) matrix built with current study’s *Desulfobacterota* class *QYQD01* and high-quality reference MAGs (>80% complete) downloaded from GTDB. MAGs that are considered same species are highlighted in red (≥95% ANI) (Jain *et al*., 2018). MAGs are labelled with the given NCBI sample/isolate name and the GenBank assembly identifier in squared brackets.

**Supplementary Figure 6.**
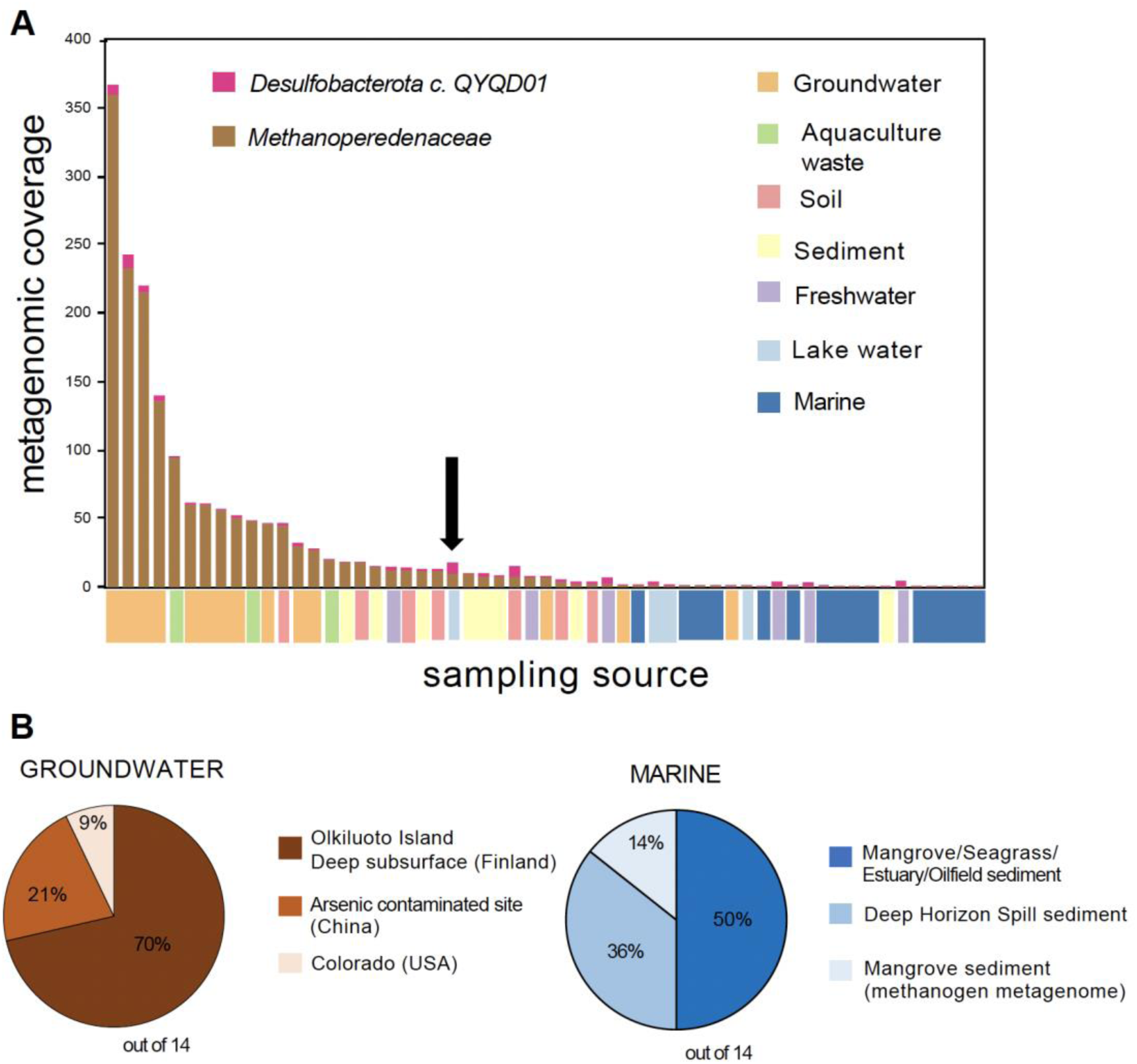
**A** Environmental distribution of co-occurring *Methanoperedenaceae* and *Desulfobacterota* class *QYQD01.* Shown are Sandpiper (via SingleM)-based SRA search results as abundance plots for *Methanoperedenaceae* and *Desulfobacterota* class *QYQD01*, based on metagenomic coverage (y-axis) across different metagenomic studies (x-axis). Note that ‘Others’ category in the pie-chart (not plotted in x-axis) harbors sites such as the iron-rich Deep Mine Microbial Observatory (DeMMO) from Casar *et al*. 2021. Metagenomic studies are arranged from left to right based on metagenomic coverage, and color-coded according to their source environment. The arrow indicates the Powell Lake (PowLake) metagenome were both metagenome assembled genomes (MAG) from ‘*Ca.* Methanoperedens’ and *Desulfobacterota* class *QYQD01* were retrieved **B** Proportions (%) of metagenomes from groundwater and marine samples, respectively, harboring both *Methanoperedenaceae* and *Desulfobacterota* class *QYQD01*.

